# A Low-Cost, 3D-Printed Biosensor for Rapid Detection of *Escherichia coli*

**DOI:** 10.1101/2022.01.04.474944

**Authors:** Samir Malhotra, Dang Song Pham, Michael P.H. Lau, Anh H. Nguyen, Hung Cao

## Abstract

Detection of bacterial pathogens is significant in the fields of food safety, medicine, public health, etc. If bacterial pathogens are not treated promptly, antimicrobial resistance is possible and can lead to morbidity and mortality. Current bacterial detection methodologies rely on laboratory-based techniques that pose limitations such as long turnaround detection times, expensive costs, inadequate accuracy, and required trained specialists. Here, we describe a cost-effective and portable 3D-printed electrochemical biosensor that facilitates rapid detection of certain *Escherichia coli* (*E. coli*) strains (DH5α, BL21, TOP10, and JM109) within 15 minutes using 500 μL of sample and costs $2.50 per test. The sensor displayed an excellent limit of detection (LOD) of 53 cfu, limit of quantification (LOQ) of 270 cfu, and showed cross-reactivity with strains BL21 and JM109 due to shared epitopes. This advantageous diagnostic device is a potential candidate for high-frequency testing at point of care as well as applicable to various fields where pathogen detection is of interest.

## 1. Introduction

*Escherichia coli* (*E. coli*) are a gram-negative bacillus that is prevalent in the gastrointestinal tracts of humans and warm-blooded animals, and well-known as one of the most significant pathogens [1]. Even though deemed harmless, numerous *E. coli* strains have acquired specific virulence attributes making them pathogenic for humans and animals. The combination of virulence factors has continued to become specific pathotypes, such as enteropathogenic *E. coli* (EPEC), shiga toxin–producing *E. coli* (STEC), enteroaggregative *E. coli* (EAEC), enterotoxigenic *E. coli* (ETEC), enteroinvasive *E. coli* (EIEC), and diffusely adherent *E. coli* (DAEC), have caused various infectious diseases in living things.

Current laboratory-based methods to detect food-borne pathogens depend on biochemical or microbiological tests, which have long turnaround times, expensive costs, and/or inadequate sensitivity and specificity [3,4]. Techniques used to analyze microbiological specimens are bacterial culturing, cytometry, immunoassays, microscopy, PCR, etc. Therefore, there is an urgent demand for a rapid and cost-effective test that is capable of identifying whole-cell bacteria with adequate sensitivity and specificity. Here, we propose a 3D-printed, host-cell detection biosensor for *E. coli* using a modified graphite pencil as a sensing electrode. This sensing platform has been used in many kinds of biosensor engineering [5]. De Lima *et al*., reported using a low-cost pencil graphite electrode coated by gold nanoparticles to detect SARS-CoV-2 on a minute scale [6]. In this study, we used thyroxine, a dual functional group molecule, and ferrocene to modify the graphite pencil electrode. This modification allows antibody conjugation and increases electrode transfer rates. Cross-reactivity studies of five different *E. coli* strains (DH5α, BL21, TOP10, and JM109) were performed to investigate the specificity of the biosensor. Due to its effective methodology, rapid diagnosis, and reduced costs, our biosensor can be a potential candidate for high-frequency testing at point of care as well as applicable to multiple fields where pathogen detection is of interest.

## 2. Materials and Methods

### 2.1. Materials

1-Ethyl-3-(3-dimethylaminopropyl)carbodiimide (EDC) with a degree of purity ≥ 98%, N-Hydroxysuccinimide (NHS) with a degree of purity ≥ 98%, gold(III) chloride trihydrate (HAuCl_4_ **·** 3H_2_O) (99.99%), sodium borohydride (NaBH_4_) with ≥98% chemical purity, cysteamine hydrochloride (Cys) with 98% chemical purity, phosphate buffer saline (PBS) solution, Silver/Silver Chloride (Ag/AgCl) paste, which was purchased from Sigma-Aldrich, was used for the reference electrode. Graphite pencil lead with 0.7 mm diameters were purchased from a local Walmart store. Anti-E.coli antibody (ab217910) was obtained from Abcam, UK. Four E. Coli-competent strains, DH5α, BL21, TOP10, and JM109, were from Real Biotech Corporation (RBC) in Taiwan. Electrochemical measurements were carried out using a CH760E potentiostat, controlled by the CH Version 12.04 software. A 3D-printed chamber containing three electrodes was used as the electrochemical cell (0.5-mL volume). Graphite pencils (0.7 mm) were used as working, reference, and counter electrodes. The graphite lead that was used as the reference electrode was coated with Ag/AgCl paste. All reagents and the deionized water (resistivity ≥18 MΩ•cm at 25 °C) used in this work were of analytical grade.

### 2.2. Electrochemical Characterization

Cyclic voltammetry (CV) and electrochemical impedance spectroscopy (EIS) were used to indicate electrochemical properties of the electrodes in each modification step according to de Lima et al., 2021 [5]. A potential window from 0.7 to −0.5 V with a scan rate of 50 mV·s-1 was set up for CV experiments. In square wave voltammetry (SWV), potentials were scanned from -0.8 to 0.4 V, corresponding to a frequency of 60 Hz, amplitude of 65 mV, and step low of 6 mV. In EIS, the frequency ranges from 0.1 Hz to 1 × 10^6^ Hz using amplitude of 10 mV and under open circuit potential was conducted. 0.1 mM KCL solution containing 5.0 mM of the mixture [Fe(CN)6]^3−/4−^ solution was used as electrolyte for all electrochemical characterization. Morphological characterizations of GPE before and after superficial functionalization were performed using the Tescan GAIA3 SEM-FIB, a specific scanning electron microscopy (SEM), from University of California, Irvine Material Research Institute (UC IMRI). SEM images were recorded with 129 to 71,000 times magnifications, acceleration voltage of 30 kV, and using In-Beam SI mode. All electrochemical experiments were carried out at room temperature (25 ± 3 °C).

### 2.3. Modification of Graphite Electrodes with Ferrocene

The working electrode (WE) polished with 2,000-grit sandpaper, and a working area of 1.5-cm length by 0.7-mm diameter was obtained. The GPE was immersed in 10 mM L-Thyroxine dissolved in dimethylsulfoxide (DMSO) for 2.0 h as a first modification step. This process allowed the graphite surface to be functionalized with iodine groups. Then, the modified graphite substrate was kept in 5 mM Ferrocene solution in toluene, which enabled ionization anchoring of Ferrocene to iodine groups of thyroxine on the electrodes. The modified electrode was washed three times in a 10 mM PBS buffer. Next, the electrodes were incubated in the solution containing 100 mM EDC and 50 mM NHS diluted in 0.1 M MES (pH = 5.0). Subsequently, 0.3 mg. mL^-1^ of anti-*E*.*coli* antibody was added into the tube containing the activated electrodes and vortex for 5 seconds. After 10 min, the antibody was immobilized onto the substrate, reached the anchoring stability, and provided a highly sensitive SWV response. In the presence of EDC and NHS, the carboxyl group of the antibodies covalently bonded to the primary amine of thyroxine. The reaction between carboxyl groups and EDC–NHS resulted in the formation of a stable ester, which undergoes nucleophilic substitution with the amine groups present on the modified electrode surface. This resulted in the formation of an amide bond between the modified GPE/Ferrocene/Thyroxine surface and the antibodies. In the final step, nonspecific sites present within the electrode surface were blocked by incubation in a 1% (mass/vol) BSA solution for 30 min.

### 2.4. Bacterial Pathogen Sensing using the Developing Sensor

For diagnosing bacterial cells, a volume of 500 μL of each bacterial strain was applied to the three-electrode configured sensing system (Fig. 1) and incubated for 5 min. After the incubation period, the sensing system was gently washed with 0.1 mM PBS buffer (pH 7.4) to remove unbound bacterial cells. Next, 500 μL of redox probe solution (5.0 mM [Fe(CN)6]^3−/4−^ in 0.1 M KCl) was injected to the chamber for voltammetry measurements and detection of current decrease due to binding of bacterial cells to the biosensor. Subsequently, the electrochemical response was monitored using the SWV technique. Specificity and cross-reactivity of the biosensor was evaluated in spiking samples. The total diagnostic time was calculated to be 13 min, which considered the sample incubation period (10 min), the time required to record two SWVs (before and after sample incubation, 2 min), and the washing step after sample incubation (1 min).

**Figure 1.**
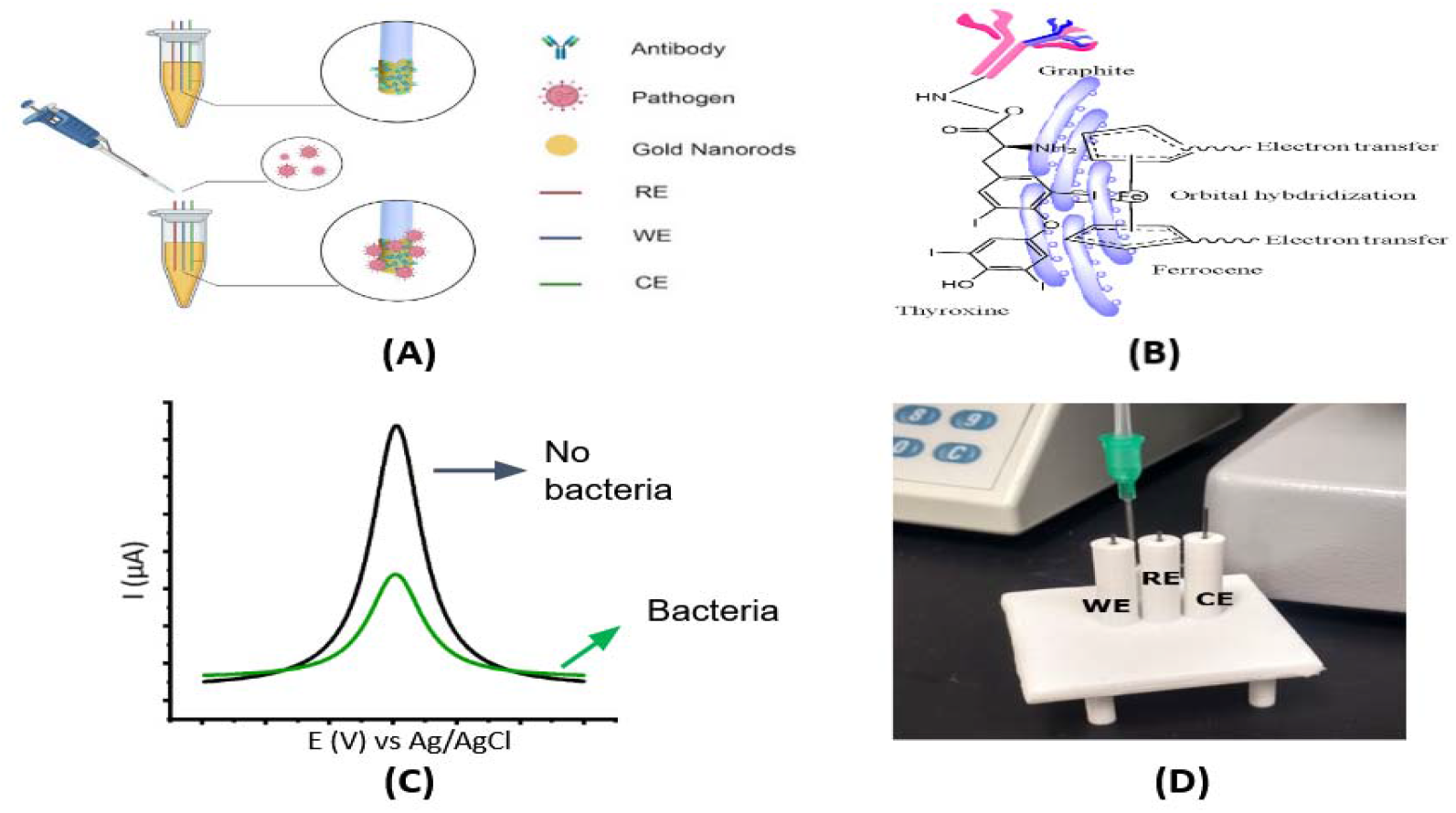
Overview of the system. **(A)** Scheme of developing an electrochemical biosensor for whole cell detection. **(B)** Thyroxine/Ferrocene functionalization on pencil graphite electrodes and antibody conjugation through ECD and NHS activation. **(C)** The concept of electrochemical response of the sensor in the presence of *E. coli* based on the current drop due to specific binding of DH5α to antibody coated onto electrodes. The *E. coli* cells bound onto electrode prevent electron transfers from the redox probe ([Fe(CN)6]^3−/4−^) to working electrodes. **(D)** An electrochemical cell containing a three-electrode configuration was made from a 3D printing protocol.

The limit of detection (LOD) and limit of quantification (LOQ) of sensor were calculated according to the four-parameter logistic curve, using Equations 1 and 2:

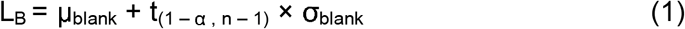

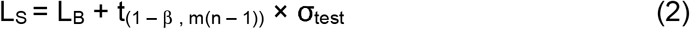

where L_B_ is a value of blank limit and L_S_ is the LOD in the signal of samples; μ_blank_ is the mean of signal intensities for n blank replicates, σ_blank_ is the standard deviation of blank replicates and σ_test_ is the pooled standard deviation of n test replicates, and t_(1 – α, n – 1)_ is the 1 − α percentile of the t-distribution given n − 1 degrees of freedom referring to the number of independent blank measurements and t_(1 – β, m(n – 1))_ is the 1 − β percentile of the t-distribution given m(n − 1) degrees of freedom referring to the number of independent sample measurements, both at 95 % confidence interval [6].

### 2.5. Specificity and Cross-Reactivity Studies

The cross-reactivity study was performed by recording the analytical signal (current decrease of the redox probe) obtained from different batches with 1000 bacterial cells on three times of measurements. The specificity of the sensors was evaluated by the analytical sensitivity value extracted from analytical curves in a concentration range from 25 to 6400 cells stored in PBS buffer (pH 7.4) and LB media. The specificity studies were carried out using the five different E. Coli-competent strains each at 100 colony-forming units (cfu). The anti-E.coli antibody was used to evaluate the capability of the sensor to detect DH5α.

The performance of the biosensor was assessed using simulated samples prepared by mixing five strains in culture media. We set a current decrease cutoff value higher than 37 μA for diagnostic purposes in accordance with the analytical response obtained for the lowest concentration of bacterial cells detected (150 cfu) in the cell–response curve. Samples that exhibited the cut off value over 15 µA were identified positive for DH5α. All sample results were analyzed and compared to those obtained from quantitative polymerase chain reaction (qPCR). The threshold cycle (Ct) values obtained by qPCR for simulated samples in which genomic DNA concentration lower than that was used in the sensing) ranged from 19.8 to 27.7.

## 3. Results and Discussion

### 3.1. Electrochemical Biosensor Design

The electrochemical device was designed to measure binding affinity between the anti-*E*.*coli* antibody and E.coli cells (**Fig. 1**). The monoclonal antibody was used as a recognition element to ensure sensitive and selective bacteria detection. The graphite that was placed in a WE was functionalized by the drop-casting method with thyroxine and ferrocene before conjugation with the monoclonal antibody (**Fig. 1A&B**). The graphite WE were polished with a 2,000-grit sandpaper to remove impurities from the surface, and a contact area of 1.5-cm length by 0.7-mm diameter was obtained. Next, to form cross-linked elements, the WE were immersed in a 10 mM L-Thyroxine in DMSO for 2.0 h as a first modification step. Thyroxine molecules have dual functional groups, such as iodine and amine groups, that facilitate the ionization and covalent attachment of ferrocene and antibodies containing carboxyl terminal moieties. Here, we leveraged thyroxine to modify the GPE’s surface with ferrocene, a versatile redox molecule. Subsequently, the coupling reaction was conducted in two different steps. First, we added the modified electrodes in a 0.1 M MES solution containing the prepared reactive intermediary EDC and NHS to form reactive coupling linkers. This enables the anchoring between the amine groups of thyroxine and carboxyl groups of antibodies, yielding Ab/thyroxine/ferrocene/GPE after 30 min of two-step activations at room temperature [7]. The electrodes were then incubated with bovine serum albumin (BSA) at 37°C for 30 min to block the electrode’s unspecific binding sites after immobilization of Ab. BSA is a common blocking protein with a high density of positive lysine residues that is commonly used for blocking procedures. Next, we exposed the sensor to samples containing bacterial cells. The decrease in the peak current of a redox probe ([Fe(CN)6] ^−3/−4^) enabled diagnosis of bacteria-free samples versus those that were infected with bacterial cells (**Fig. 1C**). Our biosensor chip is shown in Fig. 1D.

### 3.2. Characterization of the Biosensor

Electrochemical experiments were then performed to characterize the biosensor. The bare GPE presented a flat surface containing stacked carbon sheets (**Fig. 2A**). Successful formation of the ferrocene and thyroxine complex was confirmed by cyclic voltammetry, presenting a specific redox/oxidative peak at -0.15 and 0.15 e/V vs Ag/AgCl (**Fig. 2B&C**). The thyroxine coated on the WE surface after the optimized functionalization process, facilitating the subsequent anti-E.coli antibody immobilization onto the surface of the electrode. The electrochemical behavior of each functionalization step (**Fig. 2C**) was analyzed by cyclic voltammetry (CV) and electrochemical impedance spectroscopy (EIS) (**Fig. 3A**). The CV data and Bode plots revealed that the bare GPE electrode (brown line) possessed a resistance to charge transfer of 66.8 ± 8.7 Ω, indicating a small resistance toward redox conversion of ([Fe(CN)6]^−3/−4^) complex and a high rate of electron transfer on the electrode surface. This result was in agreement with the current value of 135.3 ±7.3 μA shown for the same electrode by CV plot (brown line). Next, we modified the WE with thyroxine (red line), leading to an increased current resistance to 73.1 ± 5.6 Ω and decreased current of 120.8 ± 8.9 μA. These data indicate that thyroxine acts as an electrical insulator hindering the electron transfer at the interface of the WE by inhibiting the redox probe from reaching the WE surface. Ferrocene were then anchored covalently to the surface of the GPE (blue line) through an ion bond between the iodine group from the thyroxine and the ferric groups from the ferrocene. The functionalization of ferrocene modified GPE led to decreased values of current resistance (38.7 ± 1.3 Ω) and increased transfer current (180.7 ± 2.3 μA) compared to the previous functionalization step (**Fig. 2C&3A**). **Fig. 3B** shows a magnitude view of the plots in high-frequency regions. The higher current and lower charge transfer resistance detected resulted from the greater electrocatalytic and surface area presented by the ferrocene [8], which contributed to fast electron-transfer kinetics, and thus conferring attractive features for sensor development [9]. In addition, the ammonia, free iodine, and ferric cation functional groups presented on the thyroxine/ferrocene–modified WE led to favorable electrostatic interactions of the anionic probe [Fe(CN)6]^3−/4−^,, providing an accumulation of the redox probe surrounding to the electrode interface. This improved electrochemical response, such as higher current peak, especially for whole cell detection. As a final functionalization step, we immobilized BSA to block the remaining unmodified electrochemical sites and to avoid nonspecific and undesired adsorption of other molecules (**Fig. 3C**). This step resulted in the highest resistance values (305.6 ± 17.8 Ω) and lowest current (25.4 ± 6.5 μA), suggesting a continued decline in the charge transfer kinetics after anchoring BSA due to organic materials.

**Figure 2.**
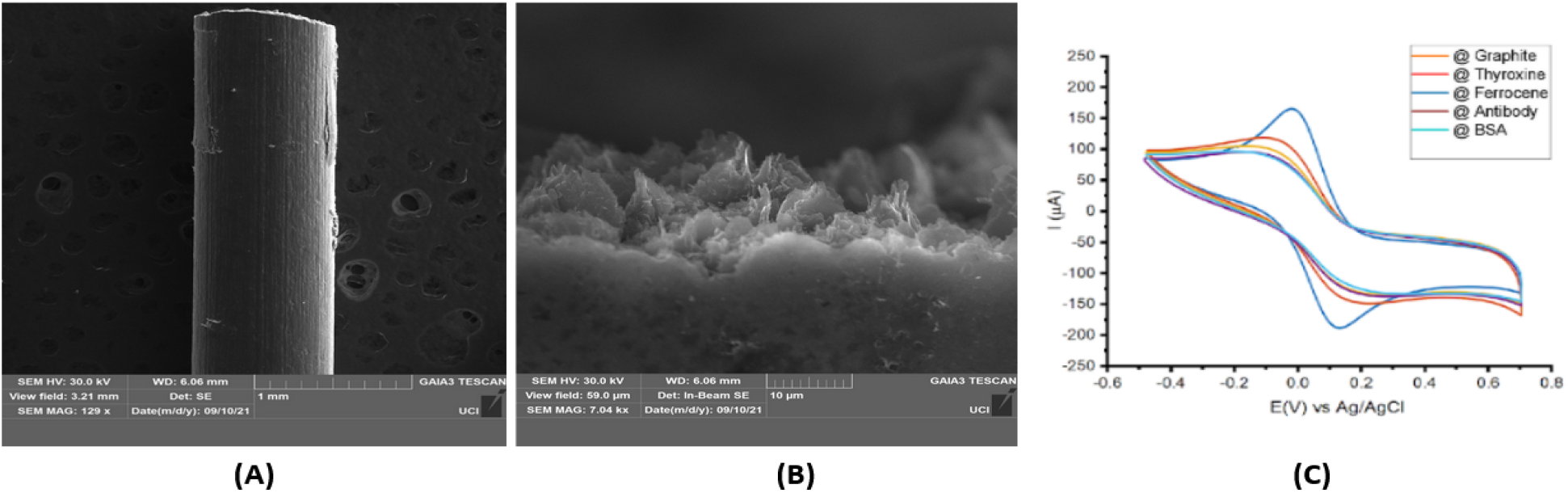
Characterization of pencil graphite electrode. (A) SEM image of bare 0.7 mm pencil graphite electrode surface at 129x magnification. (B) SEM image of 0.7 mm pencil graphite electrode after functionalization with Thyroxine and Ferrocene at 7.04kx magnification. (C) Cyclic voltammetry recorded for each bioconjugation step of the working electrode in a solution of 5.0 mM [Fe(CN)6]−3/−4 containing 10 mM KCl as the supporting electrolyte at a scan rate of 50 mV·s−1.

**Figure 3.**
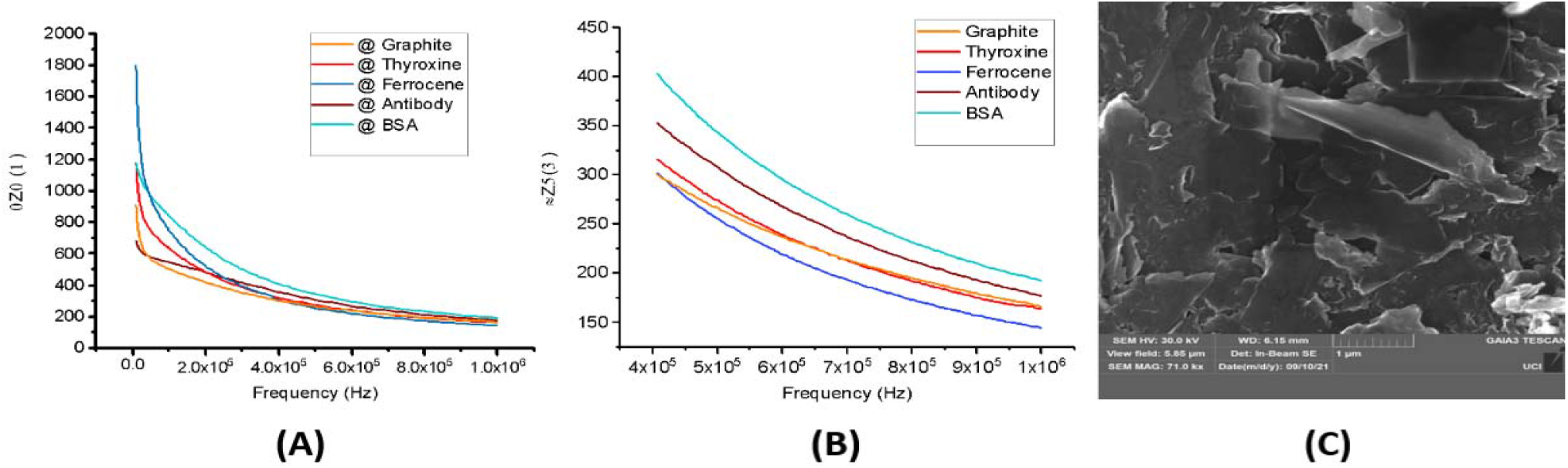
Characterization of pencil graphite electrode. **(A)** Bode plots were obtained using the same conditions as in 2C. **(B)** An amplitude view of the plots in high-frequency regions. **(C)** SEM image of working electrode after being coated and blocked by antibody and BSA, accordingly, at 71.0kx magnification.

### 3.3. Analytical Sensing of the Sensor

We used SWV for bacterial cell detection. This technique is highly sensitive, especially for detecting reversible redox species, such as potassium ferricyanide and ferrocyanide [10]. The sensing approach relied on the current signal decrease induced by specific interactions between the antibody and *E. coli* cells. The more presence of pathogen concentration leads to a more decrease in the current signal of the redox probe [Fe(CN)6]^−3/−4^. This means that bacteria bound to WE partially blocked the redox probe access WE surface. The instrumental parameters including amplitude, frequency, and step potential) were optimized to enhance capacity detection of E. coli cells. The highest peak current values for the redox probe were obtained using an amplitude potential of 90.0 mV, frequency of 115.0 Hz, and step potential of 10.0 mV). We obtained an analytical curve for different concentrations of *E. coli* in 100mM (PBS) (pH = 7.4) under optimized experimental conditions. The experiments were recorded in triplicate using an increased number of *E. coli*, from 25 to 6400 cfu. The SWV signal obtained at each cell number was shown in **Fig. 4A**. The linear regression curve was calculated at concentrations ranging from 25 to 6400 cfu of *E. coli*, resulting in an analytical sensitivity value of 55 ± 1.8 μA·cfu^-1^·mL^−1^ and a linear coefficient of 0.97 (**Fig. 4B**). Next, we conducted the optimal incubation time for detecting *E. coli* in spiking samples by evaluating the binding sensitivity parameter obtained from cell–response curves at 400 *E. coli* cells. The results were expressed as ΔI = I − I_0_, where I_0_ and I corresponds to the current recorded for the redox probe ([Fe(CN)6]^3−/4−^) before and after incubating the sample, respectively. The optimal incubation time was determined approximately in 5 to 7 min due to the highest value of the angular coefficient of the cell–response curves, referring to binding kinetics between the antibody and *E. coli*. Note that the SWV response for the redox probe [Fe(CN)6]^3−/4−^ decreased with increased concentration of *E*.*coli* due to decrease of current signal, which induced specific interaction between the *E*.*coli* and the antibody-coated WE (**Fig. 4C**). Binding of *E. coli* to the biosensor surface partially increase non-conductive layers on the electrode, leading to current drop and leading to indicate the presence of *E. coli* in the samples.

**Figure 4.**
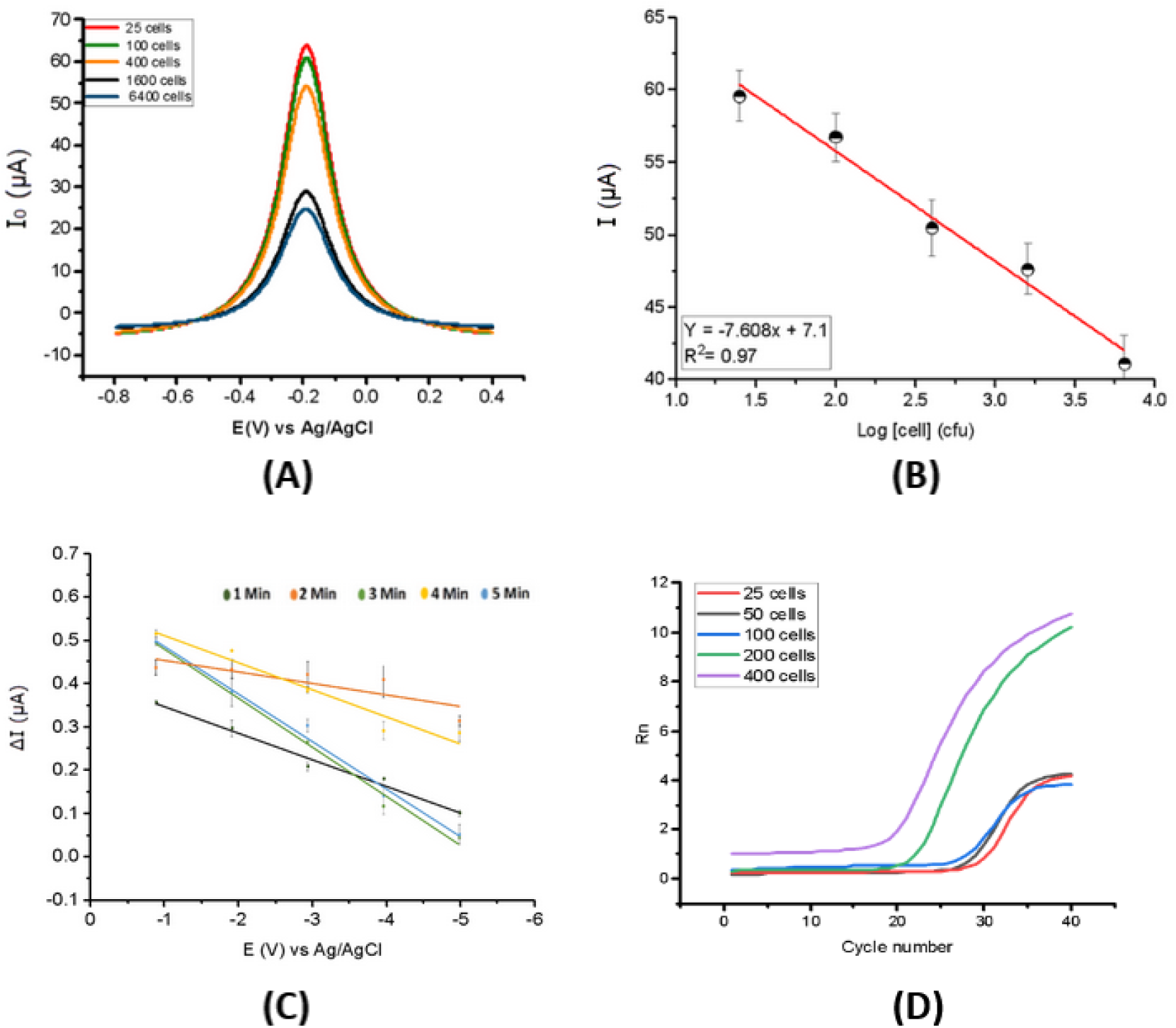
Sensing study of *E. coli* bacteria on the sensor. (**A**) Baseline-corrected SWVs of the 5.0 mM [Fe(CN)6] ^3−/4−^ redox probe containing 0.1 mol·L^−1^ KCl as the supporting electrolyte after incubating the working electrode with different concentrations of *E*.*coli* cells ranging from 25 to 6400 cells (approximately). (**B**) Linear regression (triplicate experiment) was retrieved from Fig. 4A, using the drop of current signal. (**C**) A time course (from 1 to 5 min) of binding pattern of *E. coli* cells on the modified electrode using 25 to 6400 *E*.*coli* cells. Distinct linear regression curve was obtained after 3 min. (**D**) Validation data of the presence of *E. coli* the samples by qPCR.

SWV is commonly used for assays that determine biological binding interactions and reflect the underlying binding kinetics even with small molecules [11,12]. The sensor enabled the rapid detection of *E. coli* cells (less than 100 cfu) providing high sensitivity, a low LOD in a complete 3D printing device with pencil graphite used as a WE. The threshold cycle (Ct) of the RT-PCR data (primers in **Table 1**) for cell concentration from 25 to 400 cells ranged from 21.7 to 29.3 Ct. It is important to highlight that our results (current signal) presented a high linearity (R^2^ = 0.954) with the Ct values.

**Table 1:**
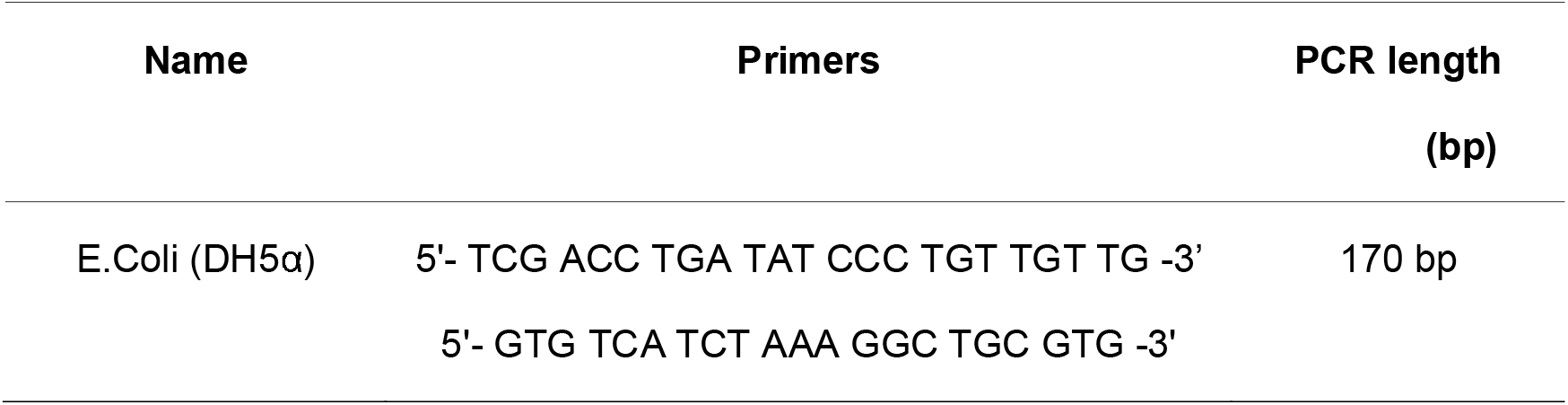
Primers used for quantitative PCR

### 3.4. Cross-Reactivity Assays

Cross-reactivity studies between *E. coli* strains were carried out to investigate the specificity of our biosensor device toward DH5α and rule out potential off-target reactivity (**Fig. 5**). Using the same experimental conditions as for DH5⍰ (**Fig. 4**), we tested four other E. coli strains including, BL21, DH5α, and JM109. DH5α (**Fig. 5A**) and TOP10 (**Fig. 5B**) showed specificity with the antibody coated into WE, which presented a current drop (ΔI) lower than the cutoff value of 37 μA obtained by SWV for the lowest *E. coli* concentration detected. BL21 (**Fig. 5C**) and JM109 (**Fig. 5D**) showed less cross reactivity to the coated antibody. The cross reactivity could be due to shared epitopes of surface proteins between *E. coli* strains that led to reduced specific binding affinity toward whole cell detection [13,14]. Development of recognition receptors for whole cell biosensors based on species-specific epitopes or aptamers should be a part of biosensor engineering.

**Figure 5.**
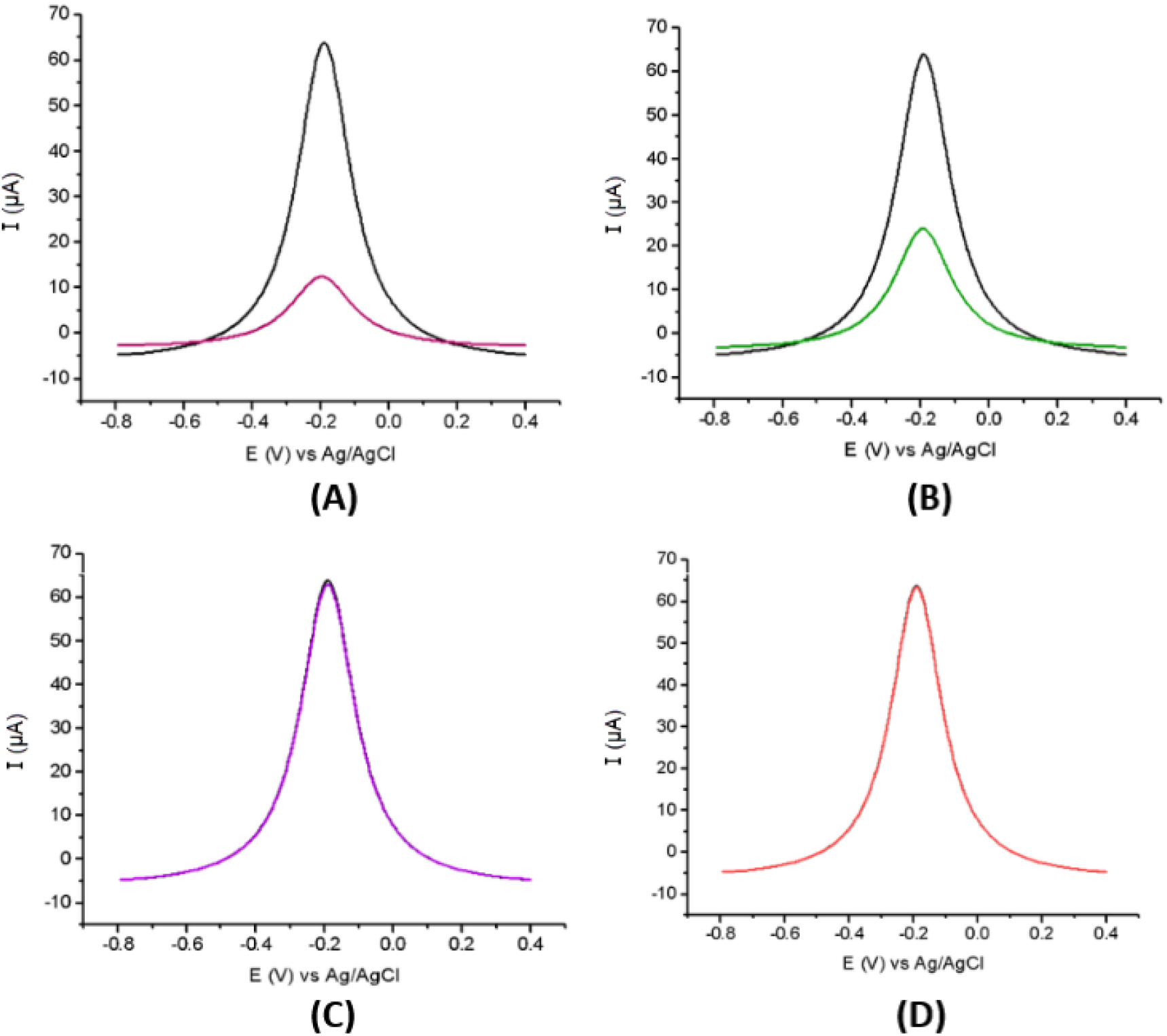
Specificity studies of the sensor using four laboratory *E. coli* strains. (**A**) DH5α, (**B**) TOP10, (**C**) BL21, (**D**) JM109. Black lines are baseline corrected SWV before challenging with bacteria sample. Experimental conditions were conducted as the same conditions described in this study.

## 4. Conclusions

We present a simple, inexpensive, and portable electrochemical biosensor based on 3D printing technique that enables diagnosis of pathogen bacteria within 15 min using 500 μL of sample and highly accessible and commercially available materials (i.e., graphite pencil leads and 3D printing), yielding a test that costs $2.50. The WE can be functionalized in less than 3 h and remains stable for over 5 d when stored in a PBS solution at 4 °C. The sensor displayed high sensitivity for detecting whole *E. coli* cell (53 cfu and 270 cfu) and showed cross-react with TOP10 and JM109 due to shared epitopes. The robustness and accuracy of the sensors were successfully evaluated by analyzing 27 spiking samples for whole cell detection, indicating that our method does not require further adaptations to accurately detect *E*.*coli* in spiking samples. Additionally, it may enable monitoring other pathogens since the modification of the electrodes with other recognition elements can be performed and easy to operate and integrate to a wireless system. Finally, the sensor can be applied for pathogens in biosafety, aquaculture, and water quality as long as the recognition elements (like aptamers and antibodies) are available.

## Author Contributions

Conceptualization

Anh H. Nguyen, Hung Cao

Data curation

Samir Malhotra, Anh H. Nguyen, Dang Song Pham,

Formal analysis

Samir Malhotra, Anh H. Nguyen, Dang Song Pham,

Investigation

Samir Malhotra, Anh H. Nguyen, Dang Song Pham,

Methodology

Samir Malhotra, Anh H. Nguyen, Dang Song Pham, Hung Cao

Writing – original draft

Samir Malhotra, Anh H. Nguyen.

Writing – review & editing

Samir Malhotra, Anh H. Nguyen, Dang Song Pham, Hung Cao

## Acknowledgement

The authors would like to acknowledge the financial support from the NSF CAREER Award #1917105 (H.C.), the NSF #1936519 (J.L. and H.C), the NIH SBIR grant #R44OD024874 (M.P.H.L and H.C.), the NIH HL107304 and HL081753 (X.X.).

